# Phenotypic Variation from Waterlogging in Multiple Perennial Ryegrass Varieties under Climate Change Conditions

**DOI:** 10.1101/2022.03.07.483244

**Authors:** Carl A. Frisk, Georgianna Xistris-Songpanya, Matthieu Osborne, Yastika Biswas, Rainer Melzer, Jon M. Yearsley

**Author notes:** RM and JMY are Joint Senior Authors.

## Abstract

Identifying how various components of climate change will influence ecosystems and vegetation subsistence will be fundamental to mitigate negative effects. Climate change-induced waterlogging is understudied in comparison to temperature and CO_2_. Grasslands are especially vulnerable through the connection with global food security, with perennial ryegrass dominating many flood-prone pasturelands in North-western Europe. We investigated the effect of long-term waterlogging on phenotypic responses of perennial ryegrass using four varieties grown in atmospherically controlled growth chambers (ambient vs 2050, +2°C and eCO_2_) during two months of peak growth. Using image analysis and PCA methodologies, we assess how multiple predictors (phenotypic, environmental, genetic and temporal) influence overall plant performance and productivity. Long-term waterlogging was found to reduce leaf-colour intensity, with younger plants having purple hues indicative of anthocyanins. Plant performance and yield was lower in waterlogged plants, with tetraploid varieties coping better than diploid ones. The climate change treatment was found to reduce colour intensities further. Flooding was found to reduce plant productivity via reductions in colour pigments and root proliferation. These effects will have negative consequences for global food security from facing extreme weather events and flooding. Our approach can be adapted as plant health diagnostics tools via remote sensing and drone-technology.

## 1. Introduction

Predicting plant responses caused by climate change is a fundamental challenge that will increasingly impact coming generations. Identifying how the various components of climate change are likely to affect plant responses will inform options of optimal mitigation strategies to minimise any negative effects (Canadell and Raupach, 2008; Pareek et al., 2020). While this is a complex issue affecting all ecosystems, it is especially important for grassland ecosystems due to the unavoidable connection with global food security through agriculture, crop lands and pasture lands (Kipling et al., 2016a; Lecerf et al., 2019; Raza et al., 2019; Tester and Langridge, 2010). There is a growing concern for the biodiversity, health, productivity and ecological transformation of grasslands from the changing climate (Goliński et al., 2018; Jentsch et al., 2009; Kipling et al., 2016b; Qi et al., 2018; Wang et al., 2018).

Grasslands are defined as ecosystems dominated by the Poaceae (grass) taxonomic family, with the ecology varying widely depending on species composition, edaphic factors, topography, management and climate. Previous studies have suggested that elevated CO_2_ levels are likely to increase photosynthetic capability in grasses, increasing Net Primary Production (NPP) and thereby total yield (Ergon et al., 2018; Yiotis et al., 2021). Meanwhile, higher temperatures are likely to extend the length of the growth season, providing longer time for sustained growth (Höglind et al., 2013; Pembleton et al., 2020). While increased ambient temperatures and elevated CO_2_ act long-term and are the main components of many climate models, altered precipitation regimes are equally important, and predicted to act both short- and long-term (Brown et al., 2019; Cullen et al., 2009; Dore, 2005). The main focus of previous grass research from altered precipitation regimes has been the effects of prolonged droughts, reduced water-availability and increased desertification (Bothe et al., 2018; Buttler et al., 2019; Cullen et al., 2014; Farfan-Vignolo and Asard, 2012; Yates et al., 2019), however many areas might see the opposite effect, leading to increased flooding (Kiely, 1999; Rosenzweig et al., 2002).

Climate change-induced alterations to precipitation regimes are expected to contribute to increased return-intervals of extreme weather events, such as severe storms and extreme flooding (Easterling et al., 2000; Semmler and Jacob, 2004). This is partly due to warmer temperatures increasing evaporation, and warmer air being able to hold more moisture, increasing the total amount of water vapour in atmospheric circulation (Hu et al., 2000; O’Gorman and Muller, 2010). Additionally, predicted changes to the precipitation seasonality are likely to change the frequency and distribution of rain events, potentially enhancing the drought-flooding dichotomy (Feng et al., 2013; Kumar, 2013).

Perennial ryegrass (*Lolium perenne*) is a common cool season pasture grass grown extensively throughout its native Eurasian-range and cultivated worldwide due to its high nutritional quality and palatability for livestock (Hannaway et al., 1999; Hunt and Easton, 1989; Minneé et al., 2019; Smit et al., 2005; Tubritt et al., 2020). Many genetically different varieties of perennial ryegrass are bred and cultivated to match the climatological conditions of specific target regions, which in turn causes differences in plant health and yield depending on local environmental suitability (Grogan and Gilliland, 2011; Helgadóttir et al., 2018). Varieties suitable in current conditions under regular precipitation regimes might prove unsuitable under periods of increased flooding (Mustroph, 2018). Flooding can cause long-term waterlogging, affecting overall plant health and total yield (McFarlane et al., 2003; Striker, 2012) and grassland ecosystem function (Fay et al., 2008). It could also impact plant morphology due to phenotypic plasticity (Mizutani and Kanaoka, 2018; Münzbergová et al., 2017). Decrease in growth due to an altered phenotypic response can have devastating impacts on global food security due to the bottom-up reliance of agricultural system productivity from grassland areas (Baldos and Hertel, 2014; Hazell and Wood, 2008). In addition, economic consequences would be especially severe for countries like Ireland and the United Kingdom with large land areas consisting of ryegrass dominated pasture lands and many rivers currently prone and predicted in the future to flood (Blöschl et al., 2019).

Here, we quantify the effects from waterlogging on perennial ryegrass performance and plant health in the light of climate change. We hypothesised that waterlogging would decrease perennial ryegrass performance and lower plant yield. We further explored whether waterlogging impacted the phenotypic plasticity of the plants. We investigated this using atmospherically controlled growth chambers and multiple commercial high-producing perennial ryegrass varieties with varying genetic backgrounds in an image analysis framework.

## 2. Materials and Methods

### 2.1. Experimental Setup

This study used four atmospherically controlled CONVIRON BDW40 walk-in growth chambers located in the PÈAC (Programme for Experimental Atmospheres and Climate) facility in Rosemount Environmental Research Station belonging to the University College Dublin in Dublin, Ireland. This facility has been used in previous studies to investigate plant responses to elevated CO_2_ (Batke et al., 2018; Yiotis et al., 2021), atmospheric paleoclimatic reconstruction (Evans-Fitz.Gerald et al., 2016; Porter et al., 2017; Yiotis et al., 2017). Two chambers were chosen to represent typical North-western European climatological conditions (CO_2_-levels at 415 ppm) while two chambers were chosen to represent predicted 2050 climate change climatological conditions with elevated CO_2_-levels (550 ppm) and a 2°C increase in temperature (IPCC, 2014). The growth chamber climatological baselines were constructed from the last thirty years of meteorological data (1989 – 2018) collated from the meteorological station located at Cork Airport and publicly accessible via the Irish Meteorological Services (Met Éireann). The entire experiment simulated conditions from May to September but only climatological conditions replicating two months of optimal pasture growth (June and July) (McHugh et al., 2020) were used to investigate perennial ryegrass responses to waterlogging (**Table 1**). Four common internationally grown varieties of perennial ryegrass were used for the experiment: *Aberchoice, Abergain, Carraig* and *Dunluce* (European Commission, 2019). The varieties vary in heading date and ploidy (**Table 2**), which allowed for an intra-species comparison to identify whether genetic factors might contribute to the response to waterlogging. At the simulated start of May each chamber was populated with 80 PVC cores (50 cm × 16 cm (⌀)) filled with John Innes No2 compost (320 cores in total for the four chambers). Each core was sealed inside a plastic bag to allow half of the cores to be waterlogged further on. The 50 cm tall cores allowed for the simulation of largely natural grassland root depth (Cougnon et al., 2017; Wedderburn et al., 2010). Each core was sown with ten seeds from one of the four varieties (DAS 0, Days after sowing) and allowed to germinate. At DAS 43, the most centrally germinated seedling was kept, and the other germinated seeds discarded. The seedlings were then cut to a height of 5 cm to simulate equal growth between all replicates. Waterlogging was initiated in half of all cores on DAS 48, with the additional watering being slowly initiated over three days to reach a stable water level 2 cm above the soil. Non-waterlogged cores continued to be watered normally. Waterlogging was actively enforced for one month and then allowed to dissipate naturally. Waterlogging experiments in grasses tend to last around 15 days (e.g. de la Cruz Jiménez et al., 2017; Liu and Jiang, 2015; Ploschuk et al., 2017), with few experiments lasting up to 30 days (McFarlane et al., 2003). The longer duration was implemented due to the increased chance (and consequences) of climate changed-induced extreme flooding events (Dore, 2005; Trenberth, 2011). Chamber treatment, waterlogging, and variety placement were stratified equally between the chambers and then randomised within chambers. The stratification allowed for equal numbers of each variety in each chamber, with half of the 80 cores per chamber being waterlogged for equal comparison for all factors.

**Table 1.**
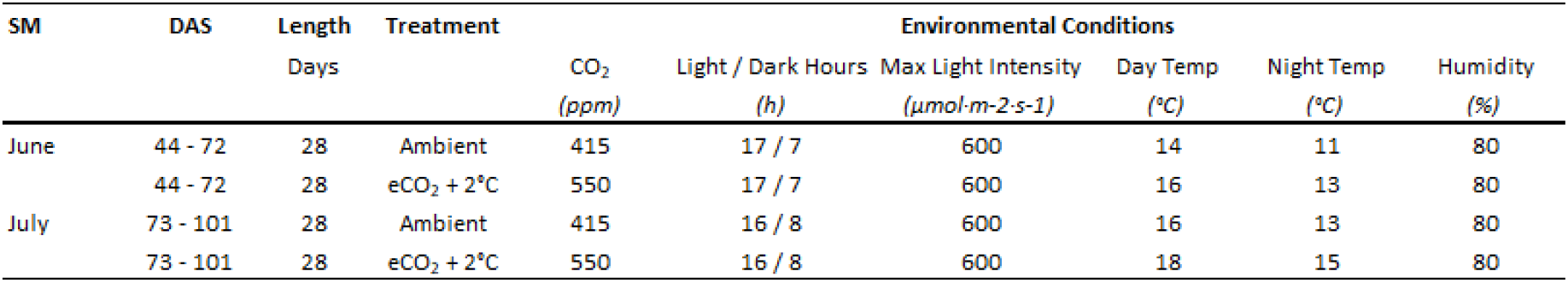
Experimental Growth Chmaber setup for the Ambient chambers (#2 and #5) and Future climate chambers (#1 and #4) simulating the months June and July in Northwestern Europe. Dawn and dusk conditions are included in the light hours. **Abbreviations**: **SM** - Simulated Months. **DAS** - Days after Sowing.

| SM | DAS | Length<br>Days | Treatment | Environmental Conditions |  |  |  |  |  |
| --- | --- | --- | --- | --- | --- | --- | --- | --- | --- |
| | | | | CO<sub>2</sub><br>(ppm) | Light / Dark Hours<br>(h) | Max Light Intensity<br>($\mu\text{mol-m}^{-2}\text{s}^{-1}$) | Day Temp<br>(°C) | Night Temp<br>(°C) | Humidity<br>(%) |
| June | 44 - 72 | 28 | Ambient | 415 | 17 / 7 | 600 | 14 | 11 | 80 |
|  | 44 - 72 | 28 | eCO<sub>2</sub> + 2°C | 550 | 17 / 7 | 600 | 16 | 13 | 80 |
| July | 73 - 101 | 28 | Ambient | 415 | 16 / 8 | 600 | 16 | 13 | 80 |
|  | 73 - 101 | 28 | eCO<sub>2</sub> + 2°C | 550 | 16 / 8 | 600 | 18 | 15 | 80 |

**Table 2.** Perennial ryegrass *(Lolium perenne)* varieties grown and water status for all chambers and variety replicates. Each water status is equally divided between all four chambers.

| Variety | Heading | Ploidity | Water Status |  |  |
| --- | --- | --- | --- | --- | --- |
|  |  |  | Logged | Normal | Total |
| Aberchoice | Late | Diploid | 40 | 40 | 80 |
| Abergain | Late | Tetraploid | 40 | 40 | 80 |
| Carraig | Intermediate | Tetraploid | 40 | 40 | 80 |
| Dunluce | Intermediate | Tetraploid | 40 | 40 | 80 |

### 2.2. Phenotypic Data Collection

Multiple sets of phenotypic data were collected each month to identify the effects on plant performance from the waterlogging. SPAD (Soil and Plant Analyzer Development) readings were conducted using a SPAD-502 Plus Chlorophyll Meter (Konica Minolta) to sample leaf chlorophyll content (e.g. Bothe et al., 2018; Dong et al., 2019). Three representative leaves were sampled from each plant, with each leaf being sampled at three places and averaged for a reliable measurement. Soil moisture was measured at 10 cm depth using a HH2 Moisture Meter with a calibrated WET sensor type WET-2 attachment (Delta-T Devices). At the end of each month (DAS 72 and 101) the maximum height of the plants was measured and plant were then harvested by cutting at 5 cm above the soil-level. The harvested material from each plant was placed on a flat white background and photographed using a high-resolution LUMIX DC-G9 camera (Panasonic) with an accompanied ColorChecker Classic chart (X-rite). This enabled the harvested material to be processed using colour-corrective image analysis techniques to identify differences in leaf colour hues (e.g. de la Cruz Jiménez et al., 2017; Hu et al., 2013; Li et al., 2014; Zhang et al., 2014). The leaves were photographed immediately after harvest to prevent structural and colour degradation (Løkke et al., 2013; Yamauchi and Watada, 2019). After the photographs all harvest material was oven dried at 65°C for one week and then weighed to measure dried biomass.

### 2.3. Image Analysis

The first stage of image analysis was to convert the colour filter array (CFA) in each RAW image from the LUMIX DC-G9 camera into a colour-corrected red-green-blue (RGB) image. This was done using a standardised pipeline with the following nine steps: 1/ subtract a black value from all pixels in the CFA, 2/ set negative pixel values to zero, 3/ divide all pixels by the maximum pixel value, 4/ correct the white balance by scaling red and blue pixels relative to green pixels in the CFA, 5/ convert pixels to unsigned 16-bit integers, demosaiced the CFA into a true-colour image using a gradient-corrected linear interpolation (Malvar et al., 2004), 6/ transform the image from the camera’s colour space to RGB, 7/ identify the 24 colours on the X-Rite ColorChecker Classic chart within the image, 8/ estimate an affine colour-transformation matrix that minimises the sum of squared deviations between the RGB colours in the image to the known colours of the 24 coloured squares, 9/ apply the colour-transformation matrix to produce a colour-correct RGB image. The camera’s image metadata was used to obtain values for black level, white-balance correction and camera colour space to RGB conversion.

The second stage of image analysis extracted RGB values from pixels that corresponded to ryegrass leaves. Ryegrass leaves were placed upon a flat, white, rectangular background which enabled the image to be cropped to the white background. The cropped image was then converted into an HSV colour-space and an initial mask created with hue values in the range 0.0 - 0.4 and 0.875 - 1.0 (corresponding to yellow, green and red hues), saturation in the range 0.2 - 1.0 and value in the range 0.0 - 0.9. Regions of the mask with connected components containing fewer than 100 pixels were removed before the mask was refined using 50 iterations of an active contours region growing algorithm and saturation values retained if they were in the range 0.25 - 1.0. The final mask was used to extract the position of pixels in the mask and their RGB values. All image processing was performed using MatLAB (version R2021a.) and its image processing toolbox (Mathworks, 2021).

### 2.4. Statistical Analyses

The ryegrass leaf RGB values from the image analysis were further processed by calculating the median value for each hue from each image of the harvested ryegrass. Median values were used due to their robust statistical properties against outliers and skewed distributions (Chen, 1998; von Hippel, 2005). To dimensionally reduce the three hues into one variable the hues were analysed using principal component analysis (PCA) from the R package *vegan* (Oksanen et al., 2020). Although variable standardisation is normally recommended (Jollife and Cadima, 2016), the hues were analysed without scaling and centring to preserve the relative values (Lever et al., 2017). We did not expect the removal of scaling to have a detrimental effect on model fitting due to the three bands having the same variances. The principal components (PC) were first tested for normality using the Shapiro-Wilks test (Shapiro and Wilk, 1965) and then analysed using Kendall’s tau rank correlation (Kendall, 1938) and Wilcoxon’s signed rank test (Wilcoxon, 1945) to test if there were any correlations and differences in mean values between the harvest groups (DAS 72 and 101) and between each water status. Kendall’s tau was used due to the higher robustness and efficiency compared to the otherwise commonly used Spearman’s rho (Croux and Dehon, 2010; Spearman, 1904). The first principal component was further used as a response variable to build a linear model to identify relevant covariates responsible for the combined hues. A similar approach has previously been used by Golzarian & Frick (2011) to investigate early growth stages of grasses. Multiple predictor variables were used to build the linear regression model: phenotypic (dried biomass, maximum height and SPAD measurements as a proxy for chlorophyll content), environmental (soil moisture as a proxy for waterlogging and chamber treatment as proxy for ambient and climate change climatological conditions), genetic (variety as a proxy for ploidy and heading date) and temporal (progression of the season and recovery from the waterlogging). The model terms were subsequently analysed using a Type II ANOVA (Langsrud, 2003; Smith and Cribbie, 2014). Model selection was then performed to analyse how removing variables in the full model would impact the AIC values and model understanding (Aho et al., 2014; Bozdogan, 1987). All statistical analyses were performed in the statistical software R (version 4.1.1.) (R Core Team, 2021).

## 3. Results

### 3.1. Plant Appearance

To estimate the effects of long-term waterlogging on perennial ryegrass we grew 320 plants in fully atmospherically controlled growth chambers, simulating typical current climatic conditions in North-western Europe and conditions as predicted in 2050 (550 ppm CO_2_ and a mean temperature increase of 2°C). Half of the plants were subjected to waterlogging 48 days after sowing (DAS) for one month. After waterlogging had begun, visible differences were observed between waterlogged and non-waterlogged plants at DAS 72 and continued to be visible at DAS 101 (**Figure 1**). The waterlogged plants by DAS 72 had stunted growth with dark brown leaf hues. The leaf morphology of the waterlogged plants also varied to the non-waterlogged plants, because leaf unfolding was disrupted, causing a concave and folded appearance of many leaves in waterlogged plants (**Figure 2**). By DAS 101 many plants had started to change in leaf colour, with many leaves possessing light green shades. This contrasts with the non-waterlogged plants that had darker green shades and a lush appearance. Leaf morphology also differed at DAS 101 between waterlogged and non-waterlogged plants, where the leaves of the waterlogged plants remained concave and light green while the non-waterlogged plants had tall, wide and lush leaves. The median true colours of the harvests at DAS 72 (**Figure 3**) and DAS 101 (**Figure 4**) showed that there was substantial variation between plants of each water treatment and chamber condition, with darkening of leaves as the growth season progressed.

**Figure 1.**
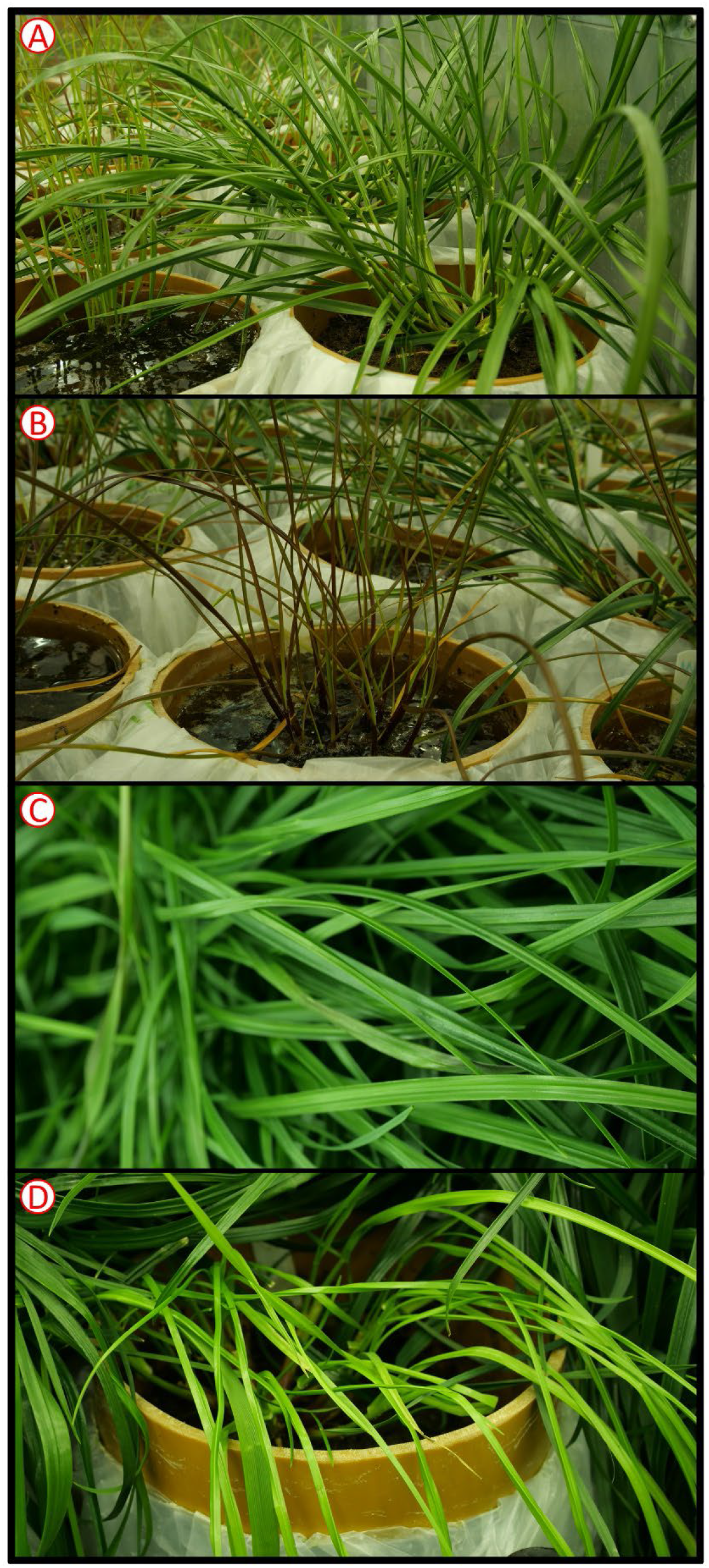
Perennial Ryegrass (*Lolium perenne*) whole plant appearance examples images for selected harvests and water status. A) DAS 72 non-waterlogged. B) DAS 72 waterlogged. C) DAS 101 non-waterlogged. D) DAS 101 waterlogged. Example images not colour-corrected.

**Figure 2.**
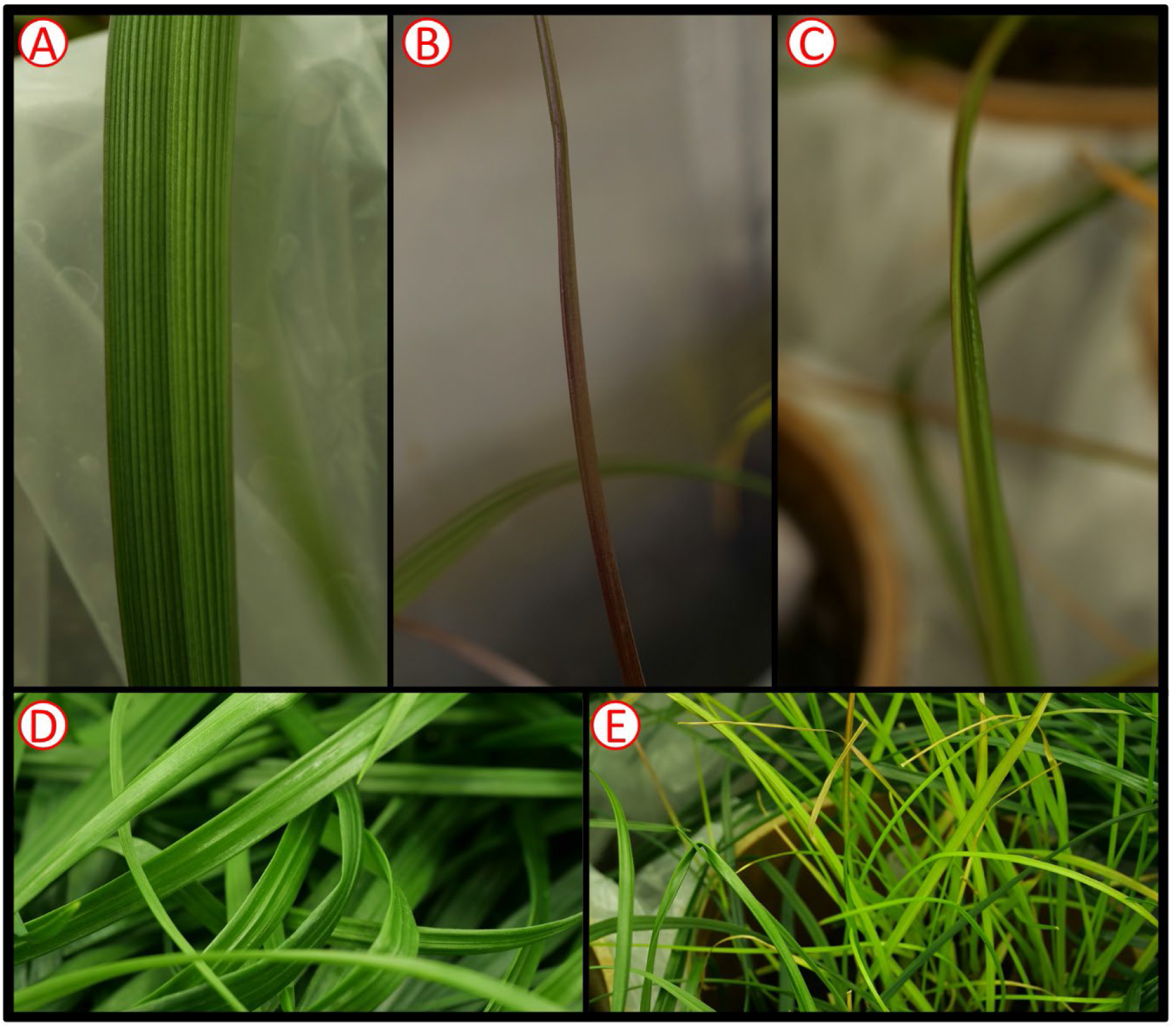
Perennial Ryegrass (*Lolium perenne*) leaf morphology example images for selected harvests and water status. A) DAS 72 non-waterlogged. B) DAS 72 waterlogged C) DAS 72 waterlogged D) DAS 101 non-waterlogged. E) DAS 101 waterlogged. Example images not colour-corrected.

**Figure 3.**
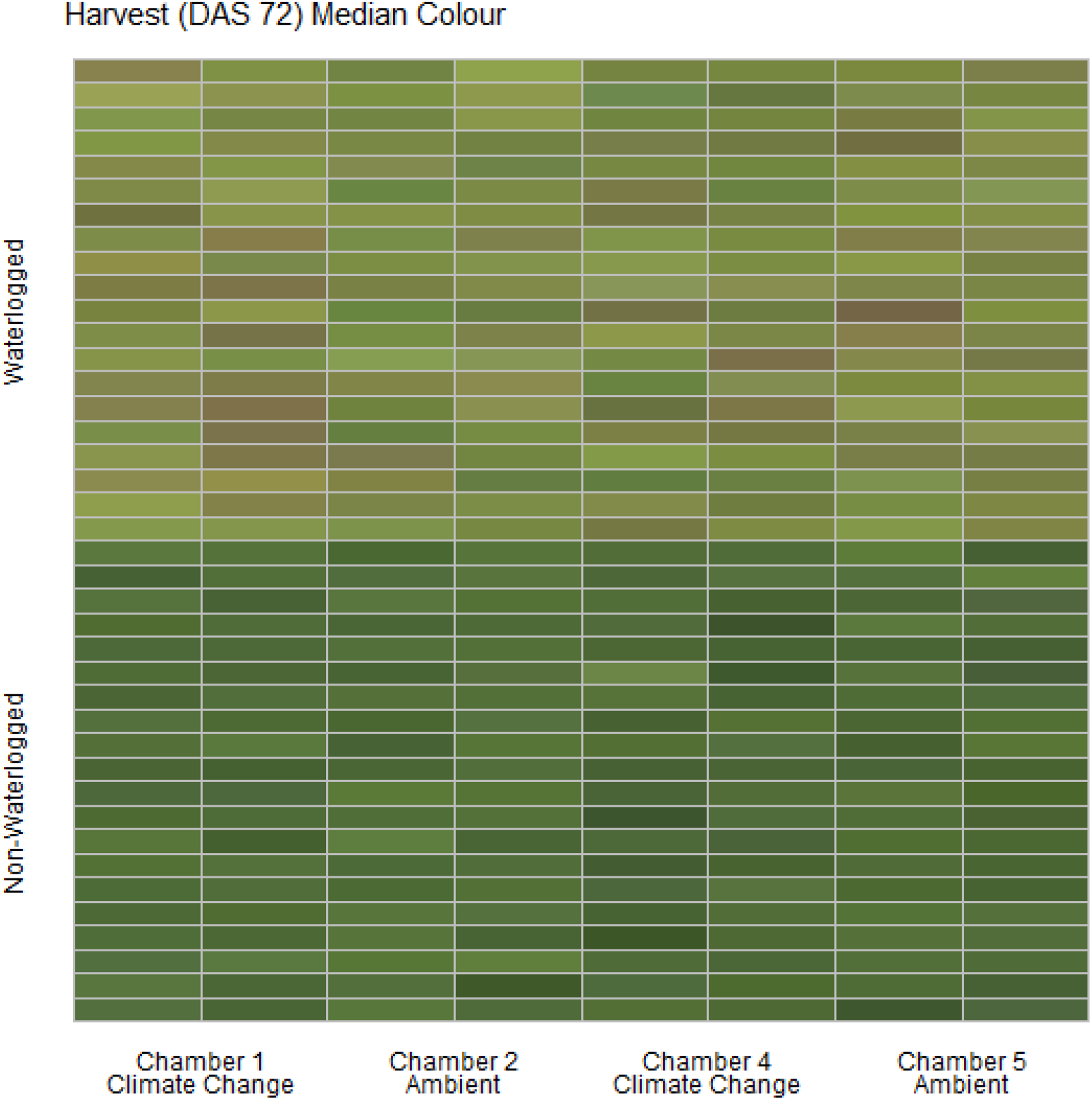
Median colour (RGB) of the harvested material for all cores on DAS 72 as identified by the image analysis, sorted on water status and chamber. Each cell represents the median true colour of the harvested material of a plant.

**Figure 4.**
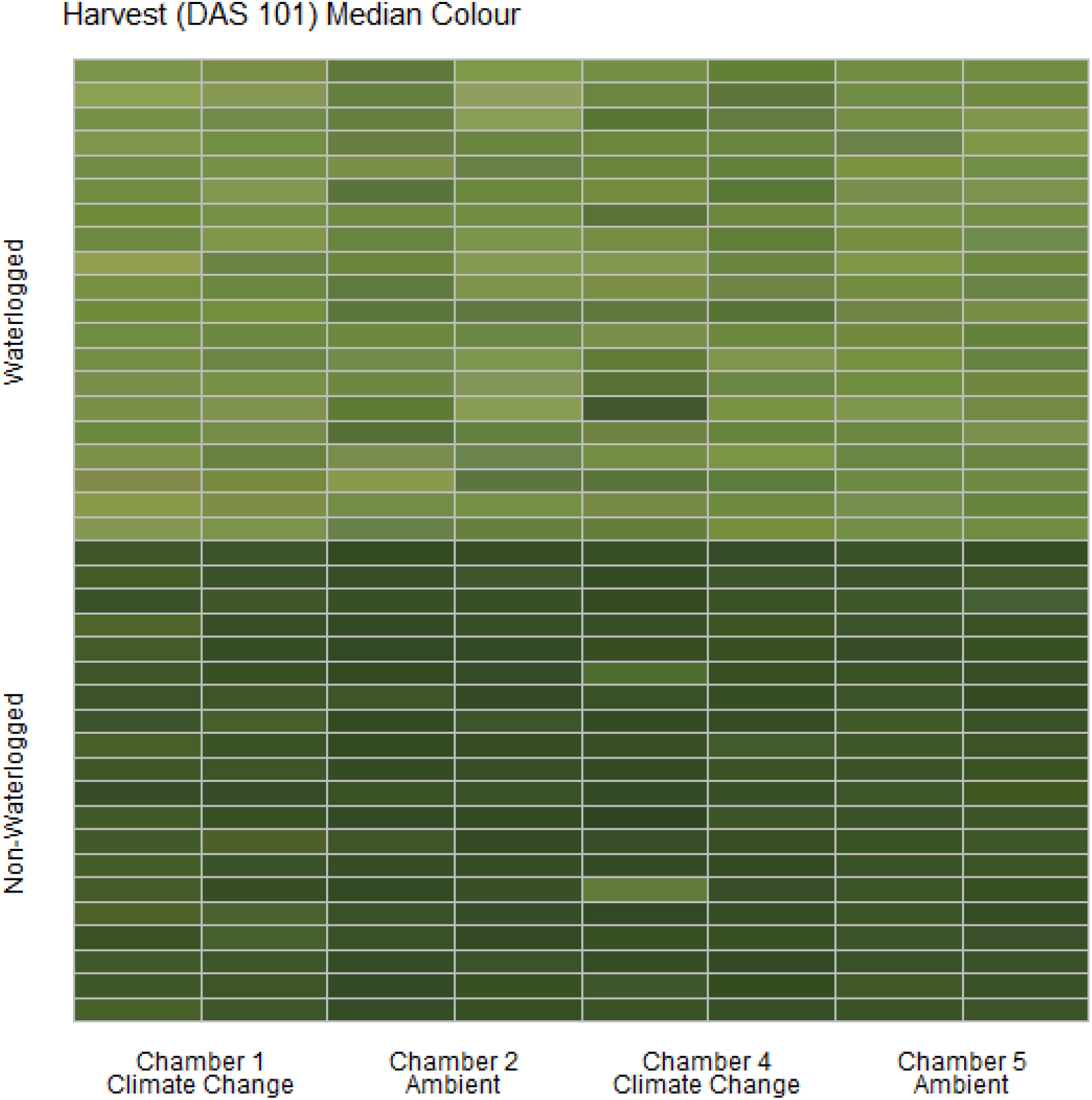
Median colour (RGB) of the harvested material for all cores on DAS 101 as identified by the image analysis, sorted on water status and chamber. Each cell represents the median true colour of the harvested material of a plant.

### 3.2. Colour Quantification Through RGB Fusion

To quantify the visible differences between the waterlogged and non-waterlogged plants, all plant leaves were harvested at DAS 72 and again at DAS 101, photographed and subjected to a colour analysis. The isolated RGB hues from the colour analysis were further modelled using a PCA-method. We hypothesised that there would be measurable colour differences within the water status, with the waterlogging contributing to lighter colours. We also expected colour differences between harvest dates, with darker colours as the growth season progressed. The PCA of the three RGB hues isolated from the harvested material from DAS 72 and 101 identified that the most variation (97.3 %) could be isolated in the first PC axis (PC1) (**Figure 5**). All three RGB hues showed a positive propensity for the first axis (**Supplementary Table 1**), illustrating that the axis describes a light beige to dark brown hue divergence (**Supplementary Figure 1**). To simplify this main leaf describing characteristics we classified the axis as overall colour intensity. The negative values of PC1 are therefore darker intensities, while the positive values are lighter intensities. The other two minor axes PC2 (1.8 %) and PC3 (0.9 %) describe pure colour hue gradients, a green-purple and an orange-blue hue divergence respectively. Kendall’s tau and Wilcoxon’s signed rank test showed that the harvested material from the waterlogged plants collected on DAS 72 were not correlated in colour to the non-waterlogged plants of the same harvest and were overall lighter (p < 0.001) and more purple (p < 0.001) (**Table 3**). The waterlogged plants on DAS 101 were positively correlated in colour intensity with the non-waterlogged plants of the same harvest while being darker (p < 0.001) and greener (p < 0.001). The harvested material from the waterlogged plants were positively correlated in colour intensity from DAS 72 to 101 with the plants becoming darker (p < 0.001) and greener (p < 0.001) as they started to recover from the waterlogging. The non-waterlogged plants were positively correlated in colour intensity between the harvests periods and became overall darker (p < 0.001) as the growth season progressed. See supplementary material for the mean differences in PC axes values between the harvests and water status groupings (**Supplementary Table 2**).

**Figure 5.**
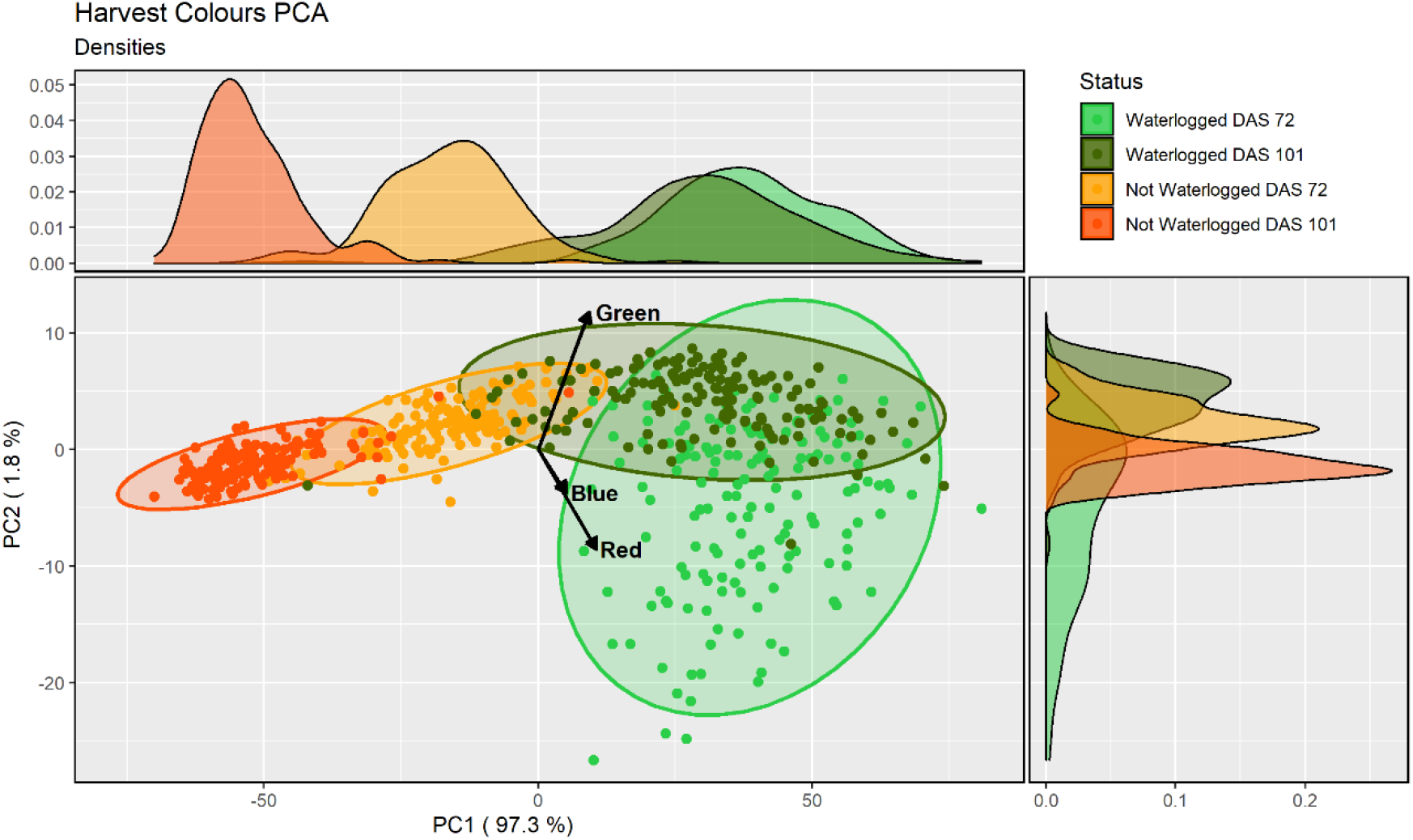
PCA-analysis of the three RGB-hues of the harvested material for all cores, water status and harvest (DAS 72 and 101) as identified by the image analysis. Each ellipse represents a 95% confidence ellipse for each group.

**Table 3.** Model statistics and significance levels for the comparison of PC axes values for the groupings harvested material (DAS 72 and 101) and water status of the cores in contrast to the other grouping. The upper part of the table compares cores within the same water status between harvests. The lower part of the table compares between water status for the same harvest. **Abbreviations**: **DAS** - Days after Sowing.

| PC Axis | Water Status |  | Harvest |  | Model Statistics |  |  |  | Wilcoxon Signed-Rank Test |  |  |
| --- | --- | --- | --- | --- | --- | --- | --- | --- | --- | --- | --- |
|  | Comparison |  | Comparison |  | Kendall's tau Rank Correlation |  |  |  | V |  |  |
|  |  |  |  |  | z | tau (t) | P - value | Significance |  | P - value | Significance |
| PC1 | Logged | | DAS 72 | DAS 101 | 9.979 | 0.532 | $< 1 \times 10^{-10}$ | *** | 1,967 | $< 1 \times 10^{-10}$ | *** |
| | Normal | | DAS 72 | DAS 101 | 2.449 | 0.131 | 0.014 | * | 36 | $< 1 \times 10^{-10}$ | *** |
| PC2 | Logged | | DAS 72 | DAS 101 | 1.757 | 0.094 | 0.079 | NS | 12,695 | $< 1 \times 10^{-10}$ | *** |
| | Normal | | DAS 72 | DAS 101 | 1.732 | 0.092 | 0.083 | NS | 445 | $< 1 \times 10^{-10}$ | *** |
| PC3 | Logged | | DAS 72 | DAS 101 | 6.007 | 0.320 | $1.893 \times 10^{-9}$ | *** | 4,232 | $1.694 \times 10^{-4}$ | *** |
| | Normal | | DAS 72 | DAS 101 | 5.495 | 0.293 | $3.907 \times 10^{-8}$ | *** | 12,875 | $< 1 \times 10^{-10}$ | *** |
| PC1 | Logged | Normal | DAS 72 | | 0.282 | 0.015 | 0.778 | NS | 0 | $< 1 \times 10^{-10}$ | *** |
| | Logged | Normal | DAS 101 | | 3.332 | 0.178 | $8.629 \times 10^{-4}$ | *** | 2 | $< 1 \times 10^{-10}$ | *** |
| PC2 | Logged | Normal | DAS 72 | | 0.285 | 0.015 | 0.776 | NS | 12,053 | $< 1 \times 10^{-10}$ | *** |
| | Logged | Normal | DAS 101 | | -0.558 | -0.030 | 0.577 | NS | 323 | $< 1 \times 10^{-10}$ | *** |
| PC3 | Logged | Normal | DAS 72 | | 1.374 | 0.073 | 0.170 | NS | 820 | $< 1 \times 10^{-10}$ | *** |
| | Logged | Normal | DAS 101 | | 0.815 | 0.043 | 0.415 | NS | 9,506 | $1.765 \times 10^{-7}$ | *** |

Significance: P < 0.001 - '\*\*\*', P < 0.01 - '\*\*', P < 0.05 - '\*', P > 0.05 - 'NS'.

### 3.3. Phenotypic Differentiation

To understand how phenotypic, environmental, genetic and temporal factors contribute to the quantified colours of the harvested leaf material a linear regression model was created and analysed. The first PC axis (PC1), identified as colour intensity, was modelled using three phenotypic variables, two environmental variables, one genetic variable and one temporal variable. The model was analysed using Type II ANOVA and AIC to see the model performance after subsequently removing each variable individually. We hypothesised that the effects from the predictor variables would be physiologically connected and cause differences in colour intensity. We expected that higher soil moisture would result in lighter colours and cause negative physiological effects on growth by lowering biomass and maximum height. We also expected that the climate change conditions would enhance growth through increased temperature and CO_2_, causing darker colours, along with darker colours as the growth season progressed, with variations between varieties caused by inherent genetic differences.

Phenotypically, harvested material from plants with lighter colour intensities had significantly lower dried biomass (F_1,630_ = 232.44, p < 0.001), significantly lower maximum height (F_1,630_ = 57.92, p < 0.001) and significantly lower SPAD values (F_1,630_ = 250.01, p < 0.001) (**Table 4**) (**Supplementary Figures 2 – 4**). For the water status, harvested material from cores with lighter colour intensities had significantly higher soil moisture (F_1,630_ = 174.67, p < 0.001) (**Supplementary Figure 5**). The harvested material from plants in chambers in predicted 2050 climate change conditions had significantly lighter intensities than harvested material from plants in ambient conditions (F_1,630_ = 19.20, p < 0.001) (**Supplementary Figure 6**).

**Table 4.**
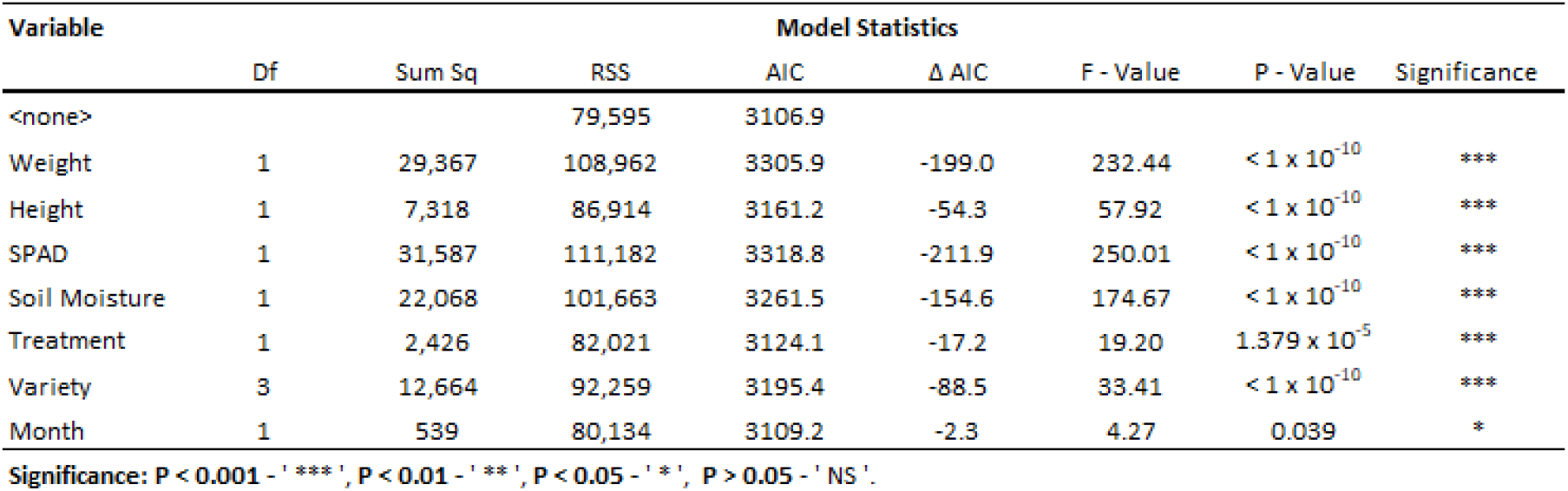
Model statistics and significance levels for the linear model in regards to the first PCA axis (Colour Intensity) isolated from the colour analysis of the harvested perennial ryegrass material. **Abbreviations**: **Df** - Degrees of Freedom, **RSS** - Residual Sum of Squares. **Model Statistics**: Anova (Type II) using the Drop1 function. **Model performance**: Adjusted R^2^ = 91.8%.

Genetically, there were significant differences in colour intensity between the varieties (F_3,630_ = 33.41, p < 0.001). *Aberchoice*, the only diploid variety, had the lightest colour intensities, while the three tetraploid varieties, *Abergain, Dunluce* and *Carraig* had darker intensities in that order, with *Carraig* having the darkest leaf colour intensities (**Supplementary Figure 7**). The harvested material became darker overall as the season progressed (F_1,630_ = 4.27, p = 0.039), with the leaves in DAS 101 being significantly darker than DAS 72. The AIC revealed that the variables had a differential importance to the model performance. SPAD values (∆AIC = −211.9), dried biomass (∆AIC = −199.0) and soil moisture (∆AIC = −154.7) were the most influential variables able to predict colour intensity, while progression of the season (∆AIC = −2.3), climate treatment (∆AIC = −17.2) and maximum height (∆AIC = −54.3) were the least influential. Overall, the linear regression model had a very high accuracy in predicting leaf colour intensity, with an adjusted R^2^ of 91.8%. See supplementary material for the linear model estimates (**Supplementary Table 3**).

## 4. Discussion

Our aim was to investigate the phenotypic effect of long-term waterlogging on perennial ryegrass. We found that long-term waterlogging reduced perennial ryegrass productivity and changed the plant phenotype. Waterlogged plants were lighter in colour than controls, with younger waterlogged plants expressing purple hues, suggestive of anthocyanins. We also found that waterlogged plants had significantly lower dried biomass and maximum height, demonstrating that waterlogging reduced overall plant performance and yield. Our study has experimentally determined the influence from waterlogging and climate change upon plant performance, with the effects from waterlogging largely overshadowing the effects from increased temperature and elevated CO_2_. The imaging methods could be developed as remote sensed diagnostics tools in combination with drone-technology to determine the influence of waterlogging on plant health in field environments.

### 4.1. Physiology and Anthocyanins

Our results identified substantial variation in leaf colouration in perennial ryegrass caused by long-term waterlogging. This variation was mainly in colour intensity, with waterlogging resulting in lighter colour intensities. Previous studies have shown that waterlogging can reduce concentrations of plant pigments, mainly chlorophyll, resulting in lighter colours and reduced photosynthetic capability (e.g. Barickman et al., 2019; Close and Davidson, 2003; Cotrozzi et al., 2021; Li et al., 2011; Ou et al., 2011; Pang et al., 2004; Simova-Stoilova et al., 2012; Smethurst and Shabala, 2003; Zhang et al., 2015). We also observed significant variation in hue, with waterlogging resulting in red/blue (purple) shades. This is especially pronounced for the waterlogged plants in DAS 72, with large variation being observed between the plants. These purple shades are likely caused by an accumulation of secondary anthocyanin metabolites, with multiple compounds having been identified in grasses (Clifford and Harborne, 1967; Fossen et al., 2002; Petrella et al., 2016). This suggests that individual plants respond differently to waterlogging by accumulating anthocyanins of varying degree. This is in agreement with previous research that has suggested that plants accumulate anthocyanins as a response to environmental stressors (Chalker-Scott, 1999), for example drought (e.g. Cirillo et al., 2021; Close and Beadle, 2003; Kovinich et al., 2015; Li et al., 2018). One of the main suggested benefits being protection against DNA damaging UV-B radiation that can reduce photosynthetic capability (Hoch et al., 2001; Rozema et al., 1997; Steyn et al., 2002; Teramura and Sullivan, 1994). Anthocyanin accumulation has also been found as a response to phosphorus deficiency (e.g. Sarker and Karmoker, 2011; Shaikh et al., 2008; Ulrychová and Sosnová, 1970). This is a likely cause of the observed purple shades in our waterlogged plants for two reasons. Firstly, phosphorus has been found to become soluble during excess water availability and migrate down the soil column (Sinaj et al., 2002) and secondly, altered plant nutrient uptake during periods of excess water abundance might make the soil nutrients (e.g., phosphorus) temporarily unavailable (Elzenga and van Veen, 2010). These effects are expected to be especially pronounced for younger plants, as we observed from our results, due to the undeveloped root system occupying only the upper part of the soil column. Additional root growth and reduction in soil moisture would allow the plants to reach the migrated phosphorus and start to recover via normalised nutrient levels, as observed with the reduction in purple hues from the waterlogging in DAS 101. Our results strengthen the consensus of recent studies that phenotypic colour variations can effectively be quantified through image analysis and analysed to detect physiological effects from waterlogging (e.g. de la Cruz Jiménez et al., 2017; Ventura et al., 2020), although detailed soil nutrient analysis would be needed to confirm our findings in connection to phosphorus availability.

### 4.2. Plant Performance

Our results showed reductions in dried biomass and height in response to long-term waterlogging for all perennial ryegrass varieties. This likely stems from lower photosynthetic capability caused by a physiological reduction in plant pigments as identified from the SPAD values and colour analysis, but also from hindrance to root development. Complex interactions between soil moisture and root proliferation patterns are likely one of the main drivers governing growth performance via soil nutrient absorption (Brugge, 1985; Cougnon et al., 2017; Medlyn et al., 2016; Setter and Belford, 1990; Xu et al., 2013). Previous studies have linked waterlogging to reductions in grass performance caused by altered root development (Malik et al., 2002; Ploschuk et al., 2017). This agrees with our findings, as we observed a clear reduction in root proliferation in the waterlogged cores (visual inspection, results not shown), with long-term waterlogging having previously been shown to reduce root mass in ryegrass (McFarlane et al., 2003). Another potential consequence is reduced root respiration caused by a reduction in soil oxygen levels, which has been shown to have multiple negative feedbacks on grass biomass accumulation and nutrients absorption (Dunbabin et al., 1997; Fukao et al., 2019; Trought and Drew, 1980).

There is also variation in waterlogging tolerance between grass species (e.g. Rubio et al., 1995; Rubio and Lavado, 1999; Xiao and David, 2019) and wheat cultivars (e.g. Ghobadi et al., 2017; Cotrozzi et al., 2021) based on local adaptation and specific genotypes. Genotype specific tolerance to waterlogging has been found between perennial ryegrass varieties (Byrne et al., 2017; Liu and Jiang, 2015). This was expanded upon by Pearson et al. (2011) that found multiple quantitative trait loci (QTLs) that code for morphological traits influencing the tolerance to waterlogging in perennial ryegrass. This corroborates the differential performance to waterlogging amongst our varieties, with the tetraploid varieties generally tolerating waterlogging better than the diploid variety. It is possible that the difference in performance is due to genotypic root development, which has been observed previously between perennial ryegrass varieties (Bonos et al., 2004; Deru et al., 2014; Wedderburn et al., 2010). Although, there is no clear connection between ploidy and stress tolerance to waterlogging or other environmental stressors (Kemesyte et al., 2017; Lee et al., 2019; Tozer et al., 2017; Yu et al., 2012). We hypothesise that a wide range of varieties of varying genotypes grown under combinations of environmental stressors coupled with detailed genomics analyses would reveal the genetic basis of stress tolerance, and potential genetic trade-offs that occur between phenotypic traits.

### 4.3. Practical Implications

We hypothesised that increases in temperature and CO_2_ (predicted 2050-levels) would enhance plant performance to climate change-induced waterlogging, but our results showed that perennial ryegrass responded with lighter leaf shades, suggesting reduced photosynthetic capability and reduced yields. This is in contrast to the theory that climate change will generally increase photosynthesis and productivity (Chen et al., 1996; Dusenge et al., 2019; Yiotis et al., 2021). The developmental period of the grasses could be relevant here, with our study investigating the performance during the first few months containing younger plants. It is uncertain if these effects are specific to developing plants, or applicable to mature ones as well. Waterlogging has previously been shown to affect grain-crops differently depending on species and development, with wheat being able to sustain growth regardless of the period of waterlogging, while barley being disproportionately affected in later development (Ploschuk et al., 2021, 2020). Daepp *et al.* (2001) showed that the effects of elevated CO_2_ will depend on the developmental stage of the ryegrass, suggesting that some stages are more sensitive to than others. For grasses in general, elevated CO_2_ has been suggested to intensify the reproductive period from increases in NPP (Kurganskiy et al., 2021). Climate change will lead to multiple changes in the environment: increased temperature and CO_2_ may have positive effects by extending the growing season for certain species while simultaneously flooding or drought may have negative effects on overall plant growth. To understand the effects of climate change we need to consider all environmental effects, their relative effect sizes and possible interactions to different developmental stages (Gray and Brady, 2016; Parmesan and Hanley, 2015; Tubiello et al., 2007; Zhou et al., 2020). In our case, the effect from waterlogging seems to largely overshadow the effects from increased temperature and elevated CO_2_.

To our knowledge this is the first study to investigate the combined effects of climate change (both temperature and CO_2_) and waterlogging experimentally in any plant. Previous studies have indicated that the interactive effects from elevated CO_2_ in isolation with waterlogging are inconclusive (Pérez-Jiménez et al., 2018; Shimono et al., 2012), with one recent modelling-study suggesting that climate change will reduce waterlogging stress in barley (Liu et al., 2021). However, increases in flooding due to climate change would cause more severe impacts than from current precipitation regimes (Iglesias et al., 2012; Mirza, 2011). The decrease in overall dried biomass for all varieties as a consequence of waterlogging predicted by our study has implications for global food security. A reduction in overall plant yield due to poor plant growth from flooding would potentially cause increases in fodder prices with follow-up consequences to all agricultural sectors and industries relying on fodder from pasture lands (Hazell and Wood, 2008; Kipling et al., 2016b; Manik et al., 2019). Early detection of waterlogging will likely be important to employ mitigation strategies that could minimise reductions in plant health and production yield in pasture lands.

We demonstrate that image analysis approaches can be used as diagnostic tools to investigate plant performance reductions caused by waterlogging. While most field monitoring would be time consuming and most satellite-based remote sensing products would be low resolution for this type of analysis other more navigable high-resolution options are available, for example drones (Bansod et al., 2017; Cracknell, 2018; Simic Milas et al., 2018). Drones have been shown to be a useful and cost-effective tool for agricultural surveying (Kulbacki et al., 2018; Puri et al., 2017; Tripicchio et al., 2015) and plant ecological investigation (Cruzan et al., 2016; Sun et al., 2021; Tay et al., 2018; Zellweger et al., 2019). The main benefits comes from the use of high-resolution multi- and hyperspectral cameras which capture a wide-range of light wavelengths used to infer plant physiological parameters (e.g. Li et al., 2020; Tao et al., 2020; Papp et al., 2021). Our results showed that differences in colour intensity could be observed between perennial ryegrass varieties, suggesting that the imaging analysis method could be developed further to identify closely-related varieties or perhaps different grass species using remote-sensed colour distributions. Recent studies have also shown the usefulness of drones to monitor the effects of waterlogging on agricultural systems (e.g. Boiarskii et al., 2019; Den Besten et al., 2021; León-Rueda et al., 2021), illustrating that our imaging approaches could be adapted to work as diagnostic tools with drone-technology. Applications using integrated monitoring of plant health will become increasingly important as climate change-induced extreme weather events become more prevalent.

## Acknowledgements

Thanks goes out to Sónia Negrão for providing assistance with the equipment acquisition. This project is funded under the EPA Research Programme 2014-2020 (Grant: 2018-CCRP-MS.52). The EPA Research Programme is a Government of Ireland initiative funded by the Department of the Environment, Climate and Communications. It is administered by the Environmental Protection Agency, which has the statutory function of coordinating and promoting environmental research. GX-S, MO and YB acknowledges funding from the Thomas Crawford Hayes Fund administered by the four NUI constituent Universities.

## Author Contributions

**CAF –** Design of the research; Performance of the research; Data analysis, collection or interpretation; Writing the manuscript. **GX-S –** Performance of the research; Data analysis, collection or interpretation. **MO –** Performance of the research; Data analysis, collection or interpretation. **YB –** Performance of the research; Data analysis, collection or interpretation. **RM -** Design of the research; Data analysis, collection or interpretation; Writing the manuscript. **JMY –** Design of the research; Data analysis, collection or interpretation; Writing the manuscript.

## Data Availability

The data supporting the findings of this study (DOI: 10.5281/zenodo.6334191) will be publicly available upon journal acceptance on the general-purpose open-access repository Zenodo developed by the European OpenAIRE program and operated by CERN.

