## Supplementary Material for "Phenotypic Variation from Waterlogging in Multiple Perennial Ryegrass Varieties under Climate Change Conditions"

**Supplementary Table 1**

Axes loadings and model variation of the Principal Component Analysis (PCA) isolated and modelled from the color analysis of the harvested perennial ryegrass material.

| **Axis** | **Loadings Color Hue** | | | **Variation** | |
|---|---|---|---|---|---|
| | Red | Green | Blue | SD | Variance (%) |
| PC1 | 0.707 | 0.621 | 0.337 | 39.222 | 97.3% |
| PC2 | -0.570 | 0.783 | -0.248 | 5.338 | 1.8% |
| PC3 | 0.419 | 0.016 | -0.908 | 3.761 | 0.9% |

**Supplementary Table 2**

Mean value and standard deviation for each grouping of water status and harvests per PC axis in the Principal Component Analysis (PCA). **Abbreviation: DAS** - Days after Sowing.

| **Water Status** | **Harvest** | **PC1 ( 97.3 % )** | | **PC2 ( 1.8 % )** | | **PC3 ( 0.9 % )** | |
|---|---|---|---|---|---|---|---|
| | | Mean | SD | Mean | SD | Mean | SD |
| Logged | DAS 72 | 38.67 | 14.11 | -4.98 | 7.22 | 1.30 | 3.57 |
| | DAS 101 | 29.92 | 17.96 | 4.07 | 2.27 | 0.04 | 3.74 |
| Normal | DAS 72 | -16.27 | 11.66 | 2.19 | 2.10 | -3.33 | 2.83 |
| | DAS 101 | -52.32 | 9.90 | -1.28 | 1.57 | 1.99 | 2.29 |

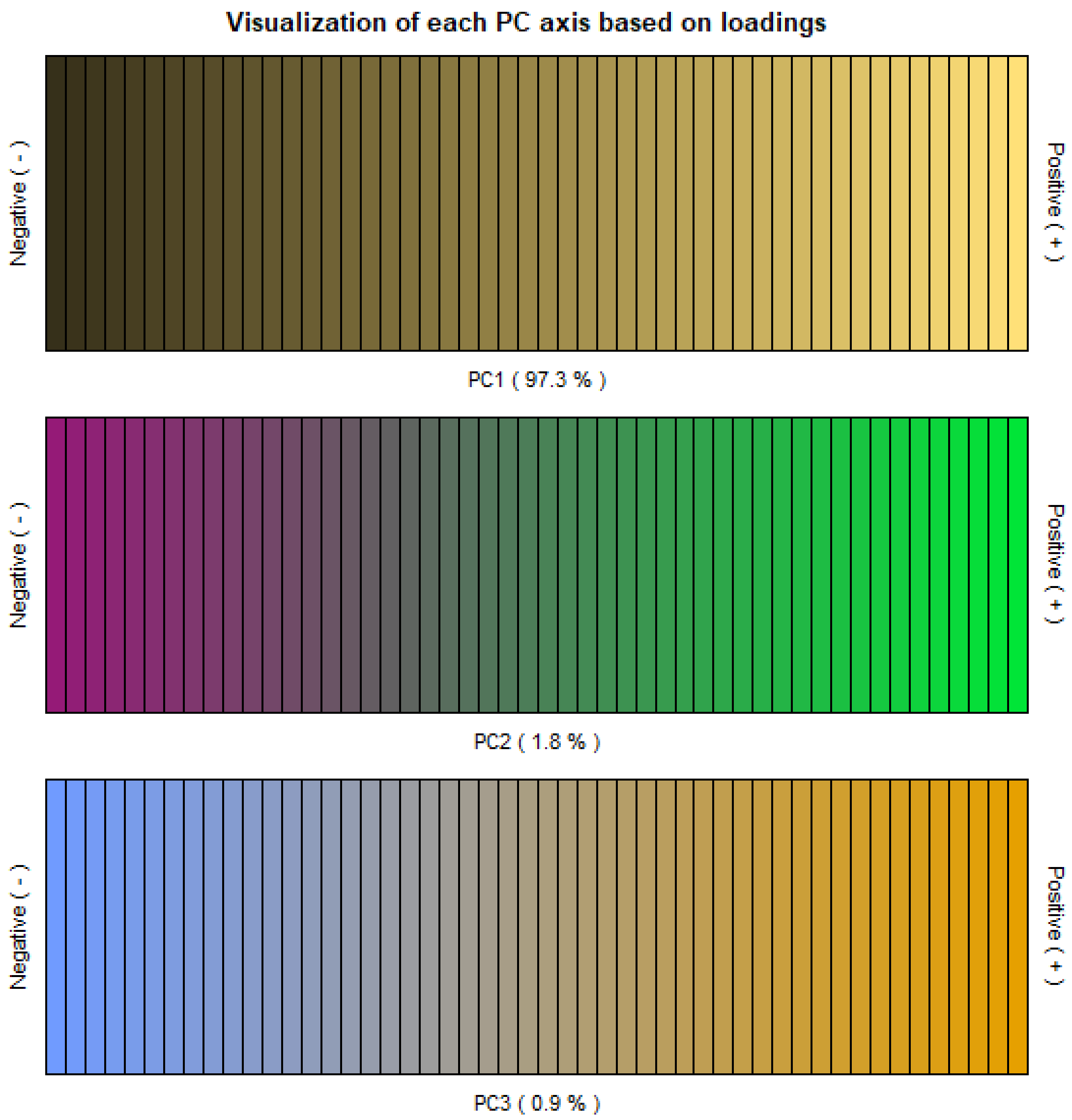


**Supplementary Figure 1.** Visualisation of the colour spectrum of each PC axis based on the loadings from each RGB hue from the image analysis.

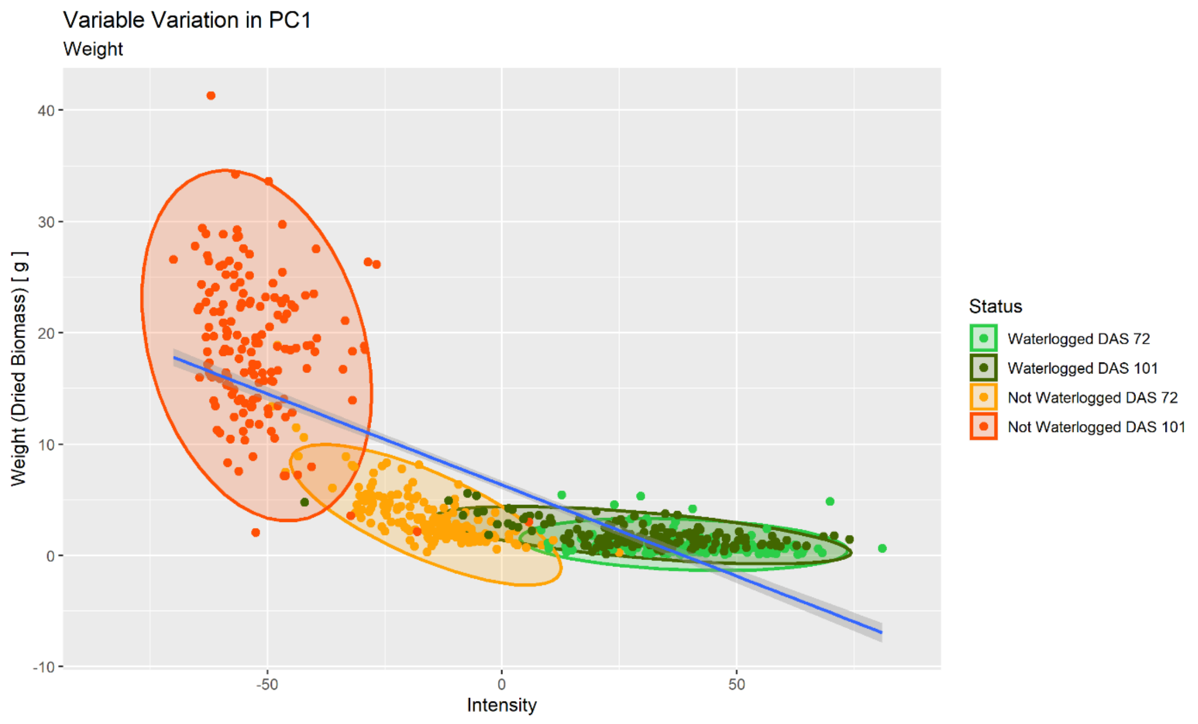


**Supplementary Figure 2.** Relationship between dried biomass and colour intensity (PC1) for all harvested cores, water status and harvest (DAS 72 and 101) as identified by the image analysis. Each ellipse represents a 95% confidence ellipse for each group. The trend line is a smoother based on linear regression (y~x) with 95% confidence intervals.

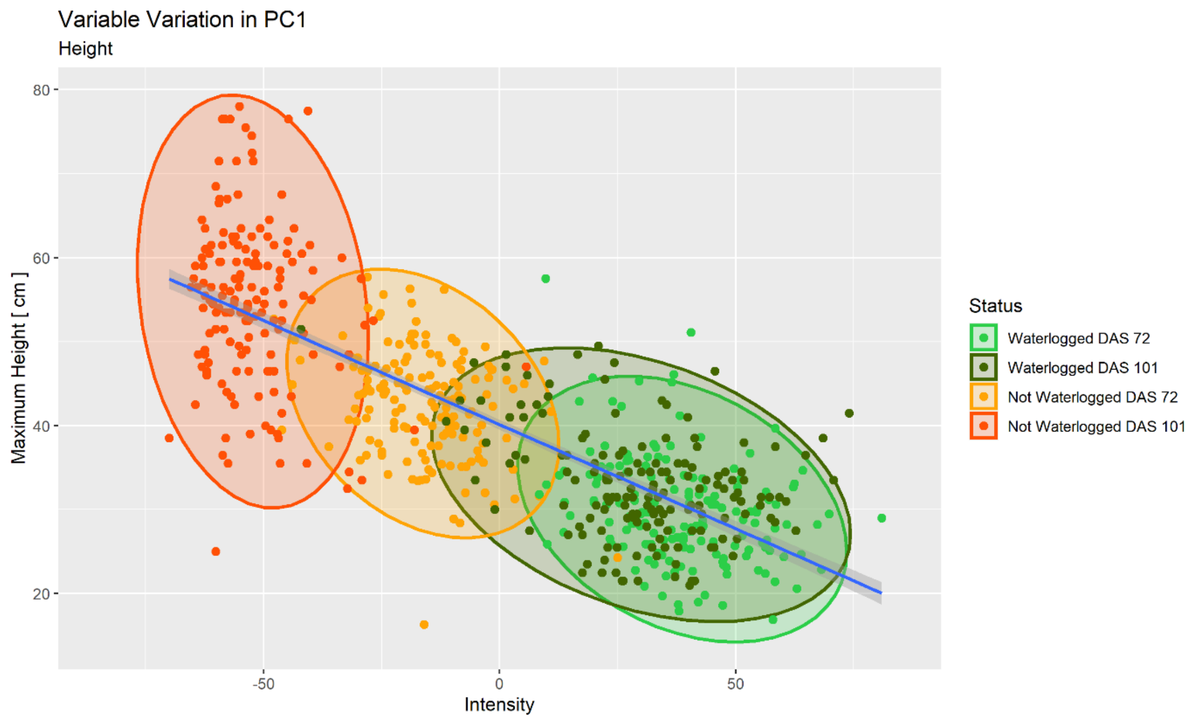


**Supplementary Figure 3.** Relationship between maximum height and colour intensity (PC1) for all harvested cores, water status and harvest (DAS 72 and 101) as identified by the image analysis. Each ellipse represents a 95% confidence ellipse for each group. The trend line is a smoother based on linear regression (y~x) with 95% confidence intervals.

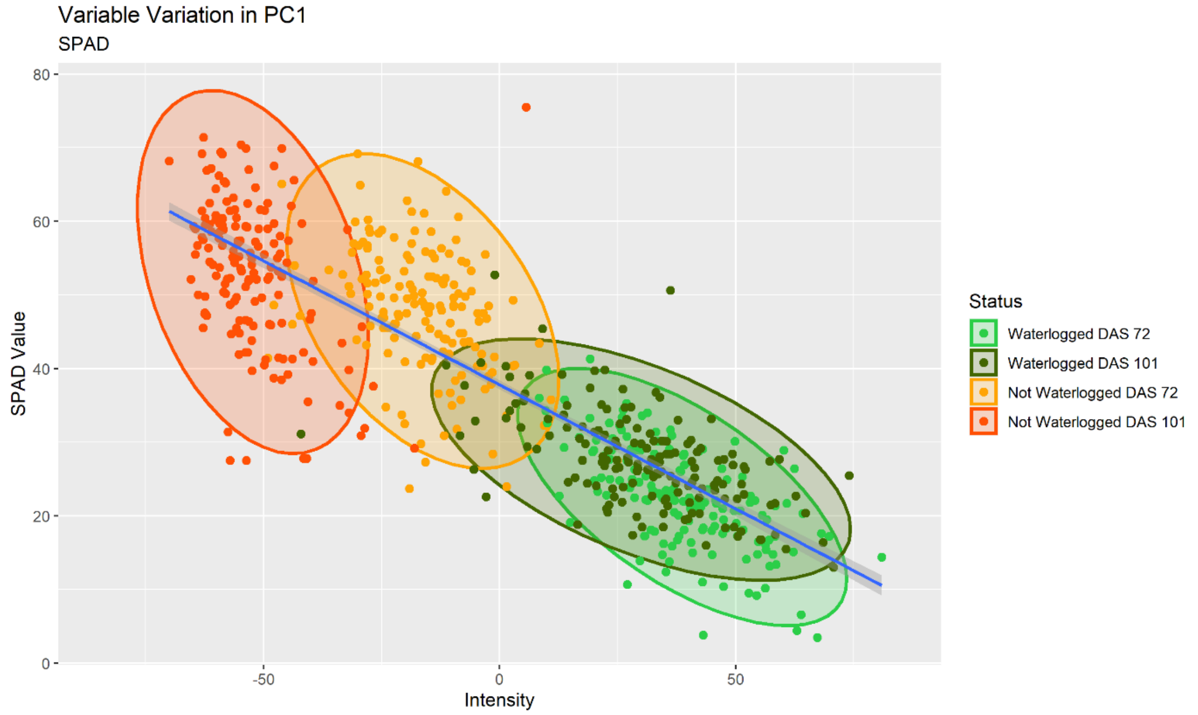


**Supplementary Figure 4.** Relationship between SPAD values and colour intensity (PC1) for all harvested cores, water status and harvest (DAS 72 and 101) as identified by the image analysis. Each ellipse represents a 95% confidence ellipse for each group. The trend line is a smoother based on linear regression (y~x) with 95% confidence intervals.

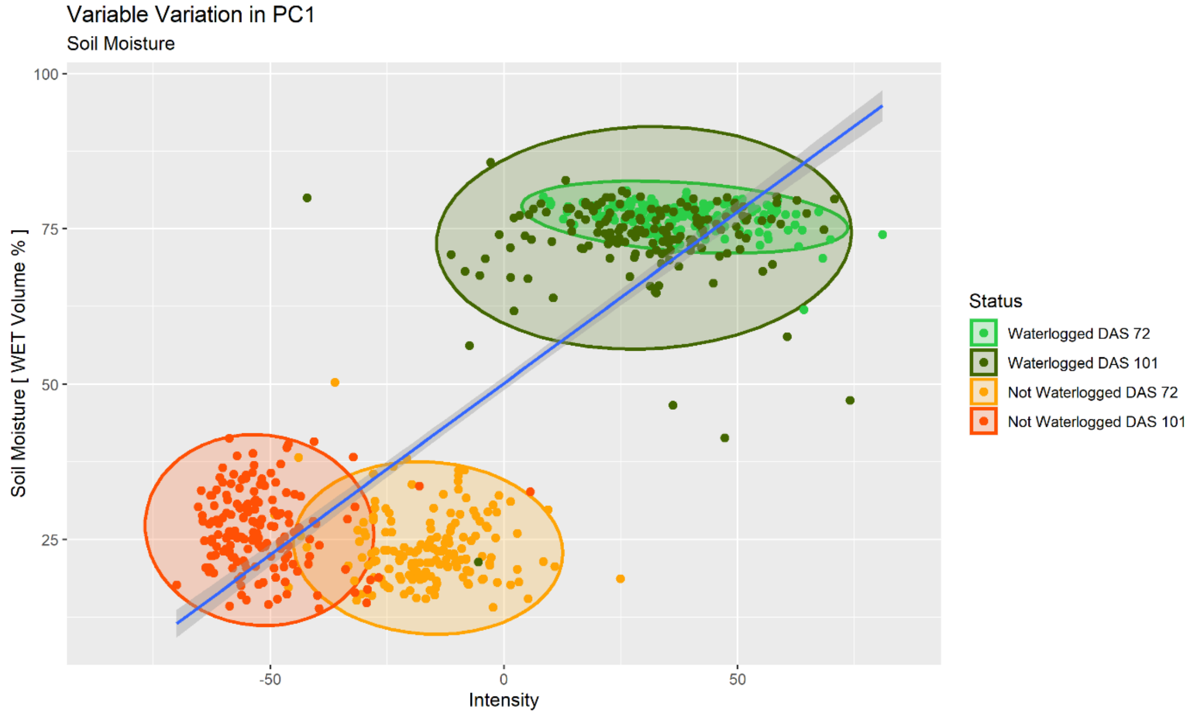


**Supplementary Figure 5.** Relationship between soil moisture and colour intensity (PC1) for all harvested cores, water status and harvest (DAS 72 and 101) as identified by the image analysis. Each ellipse represents a 95% confidence ellipse for each group. The trend line is a smoother based on linear regression (y~x) with 95% confidence intervals.

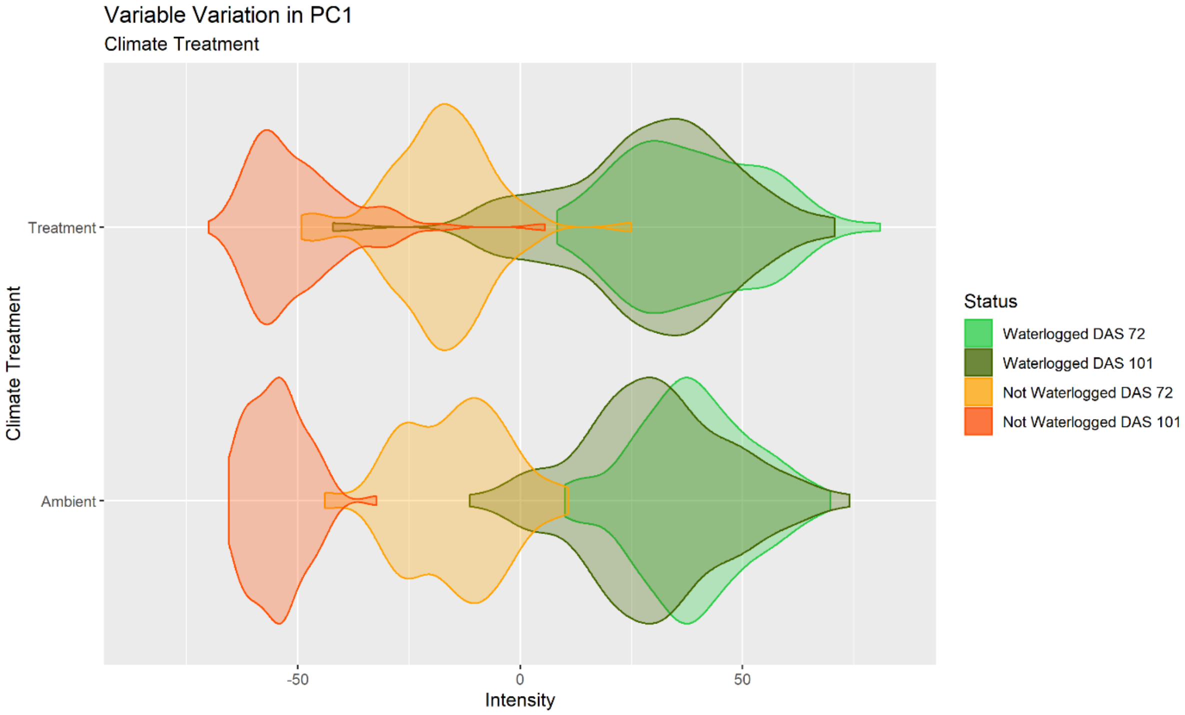


**Supplementary Figure 6.** Violin graphs describing the relationship between climate treatment and colour intensity (PC1) for all harvested cores, water status and harvest (DAS 72 and 101) as identified by the image analysis.

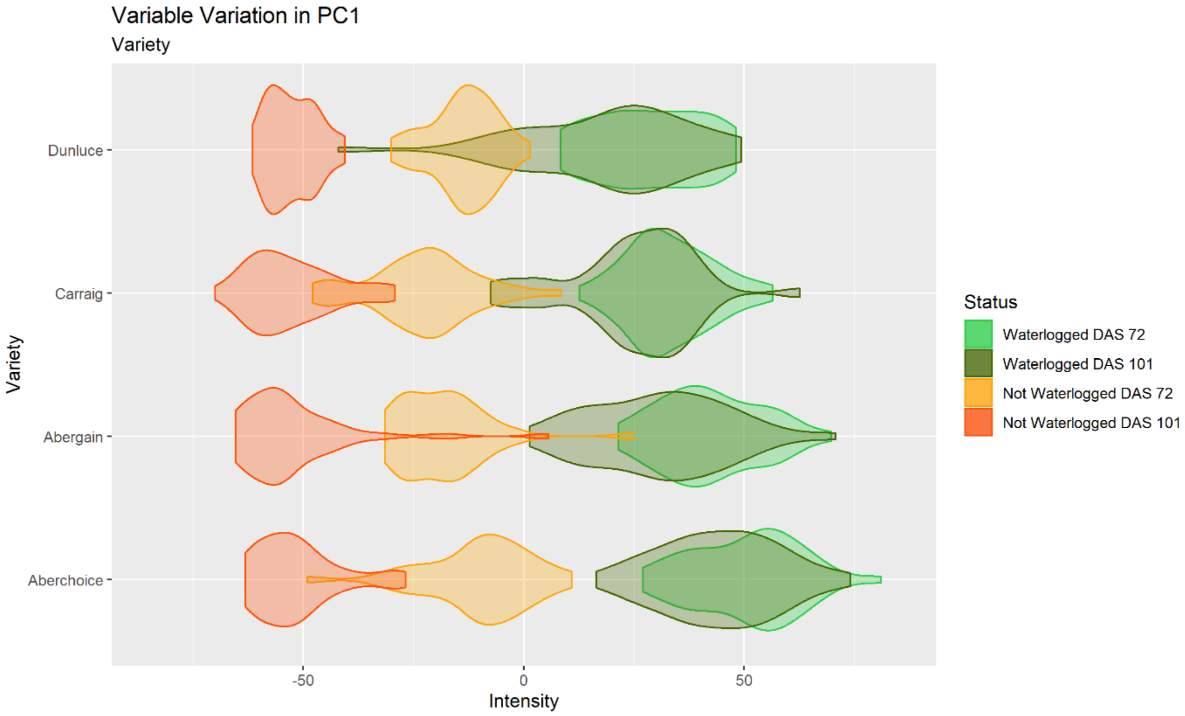


**Supplementary Figure 7.** Violin graphs describing the relationship between varieties and colour intensity (PC1) for all harvested cores, water status and harvest (DAS 72 and 101) as identified by the image analysis.

**Supplementary Table 3**

Model statistics for the variables in the linear regression model in regard to the first PCA axis.

**Intercept: Treatment** - Ambient, **Variety** - Aberchoice, **Harvest** - DAS 101. Extension of **Table 4.**

**Abbreviation: DAS** - Days after Sowing

| Variable | | Model Statistics | | |
|---|---|---|---|---|
| Main variable | Sub-factor | Estimate | Std. Error | t - value |
| Intercept | | 42.540 | 4.385 | 9.701 |
| Weight | | -1.481 | 0.097 | -15.246 |
| Height | | -0.483 | 0.063 | -7.611 |
| SPAD | | -0.850 | 0.054 | -15.812 |
| Soil Moisture | | 0.446 | 0.034 | 13.216 |
| Treatment | $eCO_2$ + 2°C | 3.980 | 0.908 | 4.382 |
| Variety | Abergain | -6.793 | 1.273 | -5.336 |
| | Carraig | -11.429 | 1.288 | -8.877 |
| | Dunluce | -10.805 | 1.297 | -8.329 |
| Harvest | DAS 72 | 2.377 | 1.151 | 2.065 |
